# Endothelia extrude apoptotic cells to maintain a constant barrier

**DOI:** 10.1101/268946

**Authors:** Tara M. Mleynek, Michael Redd, Aubrey Chan, Yapeng Gu, Dean Y Li, Jody Rosenblatt

## Abstract

The vascular system is lined with endothelial cells that, although only existing in a single monolayer, are key in the regulation of vascular barrier function. One of the major challenges these cells face is a routine exposure to environmental stressors that can induce apoptosis. Uncontrolled apoptosis in the endothelial monolayer threatens the ability of the cells to maintain their barrier function, resulting in vascular dysfunction. Therefore, we sought to identify ways in which endothelia maintain a cohesive monolayer during apoptotic events. We found that endothelial cells fated die will undergo a process of apoptotic cellular extrusion, similar to what has been described in the epithelium. We further show that endothelial extrusion uses a conserved S1P-S1PR2-RhoA signaling pathway in order to induce the formation of an actin ring that contracts closed, forcing the dying cell out of the monolayer while simultaneously filling in the gap left behind. Thus, endothelial extrusion successfully removes an apoptotic cell before it compromises the monolayer, preserving the barrier function.

## Introduction

The luminal surface of blood vessels is lined with a monolayer of interconnected endothelial cells that provide a semi-selective barrier between the intravascular compartment and the surrounding tissues (Udan *et al.*, 2013). Disruption of the endothelium can lead to a disruption of the barrier resulting in vascular dysfunction and misregulation of thrombosis (Yau *et al.*, 2015; Alexander *et al.*, 2009).

Similarly, epithelial cells cooperate to form a monolayer barrier for the many organs they encase. We previously identified and characterized a process termed ‘cell extrusion’ that enables epithelia to maintain a constant barrier even upon death of up to 40% of their component cells (Rosenblatt *et al.*, 2001). Extrusion is initiated when a dying epithelial cell fated to die emits the lipid Sphingosine 1-Phosphate (S1P), which binds the S1P receptor 2 (S1P_2_) on neighboring cells. S1P_2_ activation triggers Rho-mediated assembly and contraction of actin and myosin II fibers to physically force a dying cell from the epithelium and simultaneously prevent gap formation at the exit point. Apoptotic stimuli have been shown to trigger this response, preventing the subsequent loss of membrane integrity (Eisenhoffer *et al.*, 2012; Andrade *et al.*, 2011). Disruption of S1P-S1P_2_-RhoA signaling inhibits extrusion and results in poor barrier function, neoplasms, and cell invasion, indicating a critical role for extrusion in maintaining barrier integrity (Marshall *et al.*, 2011; Slattum *et al.*, 2009; Gu *et al.*, 2011).

Here we demonstrate that endothelial cells, both *in vivo* and *in vitro*, extrude apoptotic cells via S1P-S1P_2_-RhoA to maintain a constant barrier. Identification and characterization of this process not only indicates that epithelia and endothelia share a conserved signaling mechanism to protect barrier integrity but suggests new etiologies for vascular disease.

## Results

### Endothelia extrude cells fated to die in zebrafish

Endothelial cells must act collectively together to provide a protective and selective barrier for blood as it travels throughout the body. Should this barrier be breached by cell death, the force of blood pumping throughout the vasculature could cause hemorrhaging, inappropriate signaling to regions within the body, and possibly oxidative stress and additional cell death in regions of hemorrhage. Thus, we predict that like epithelia, endothelia possess a mechanism that prevents barrier disruption and hemorrhage when endothelial cells die.

To investigate this in a vascularized model organism where we could image endothelia live, we used developing zebrafish embryos. These embryos develop outside the mother and their transparency enables single cells to be imaged live in developing animals. Moreover, the transgenic kdrl:GFP zebrafish line specifically labels endothelial cells fluorescently, which allows us to directly follow endothelial cells. We focused on the zebrafish common cardinal veins (CCVs), which develop as large flat endothelial sheets between 24-60 hours post fertilization (Helker *et al.*, 2013). These endothelia provided us with an excellent model system to readily image individual cells in a live organism. (Figure 1A). Because the intrinsic programmed cell death rate is exceedingly small in developing zebrafish CCV endothelia, we triggered apoptosis using a 920nm two-photon laser directed at specific cells at low levels that would not cause completely ablate the cells (Figure 1B). Laser-treated cells typically die by apoptosis within 4-8 hours following laser treatment, as indicated by cell shrinkage and membrane blebbing (Ziegler *et al.*, 2004). As these cells die, we found that like epithelial cells, they also extrude. The extruding cells pop out of the plane while the neighboring cells close around and below them, thereby, preventing any gaps from forming as they leave (Figure 1C, Movie S1). Although apoptosis was variable between individual zebrafish embryos, one or more extrusion events could be observed in most samples.

**Figure 1:**
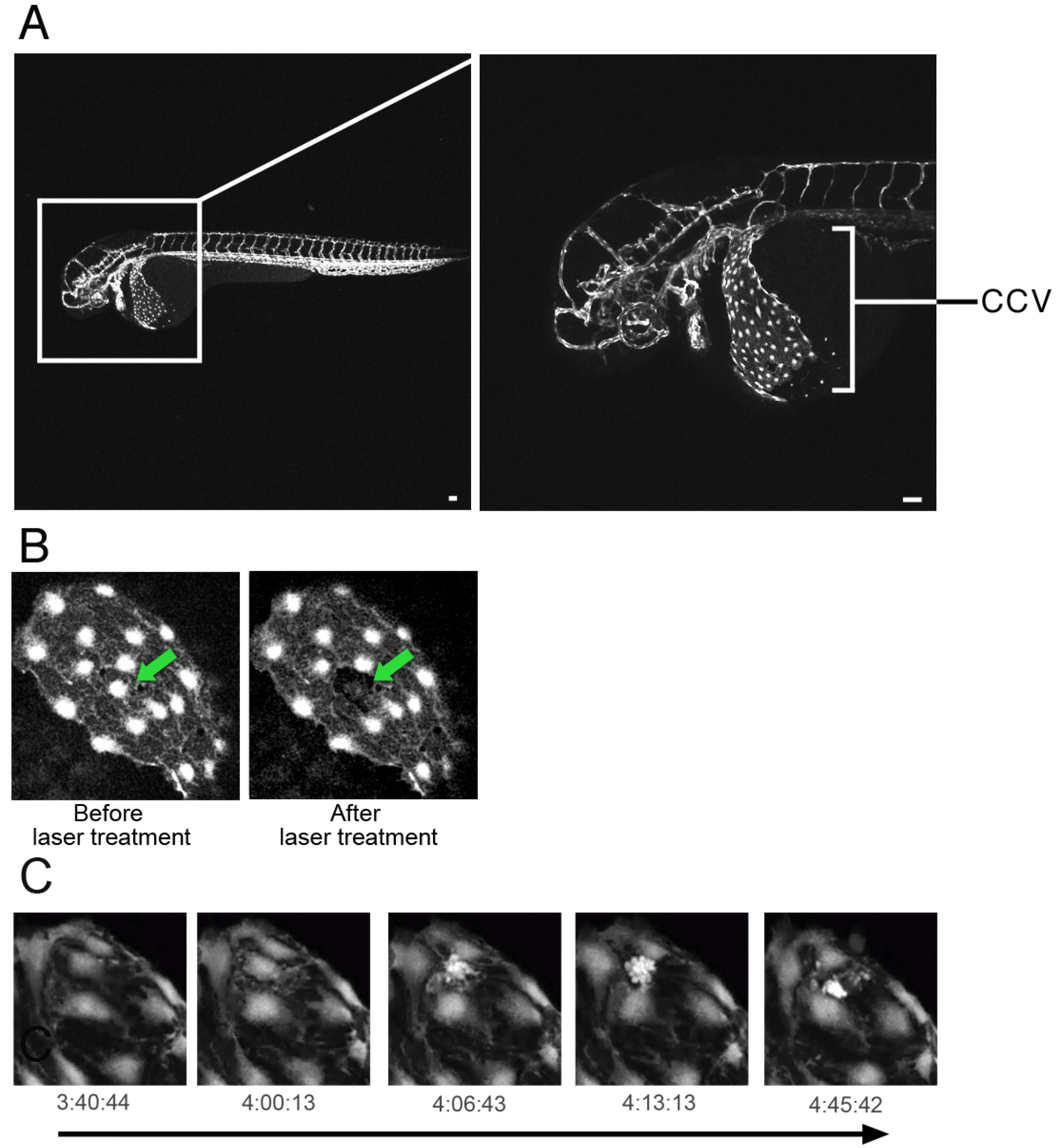
Endothelia extrude dying cells *in vivo*. A) Zebrafish with GFP labeled vasculature, inset highlights the location and morphology of the developing CCV. B) Endothelial cells in the CCV before and after laser targeting. Arrow indicates targeted cell. C) Stills from an *in vivo* movie (Movie S1) with time passage indicated below.

### Endothelia extrude apoptotic cells

We next sought to identify whether endothelial cell extrusion required the same signaling and mechanism we had previously identified for epithelial cell extrusion. Because we can more readily analyze and manipulate function in cultured cells, we developed our assay in the Human Umbilical Vein Endothelial Cell (HUVEC) line. To test if HUVEC cells targeted to die could be extruded, we grew them to confluence on fibronectin-coated glass coverslips, exposed them to UV^254^ light for approximately 45 seconds to induce apoptosis, and then immunostained for actin, DNA, and active-caspase-3. This dose of UV induced approximately 7-23% of the cells to undergo apoptosis within two hours. Yet, even with this high rate of apoptosis, we observed intact endothelial monolayers with caspase-3-positive apoptotic cells extruded, as indicated by an actin ring around and below the apoptotic cell (Figure 2A-F). To film extruding HUVEC cells, we expressed GFP-LifeAct, a protein that labels filamentous actin and filmed monolayers following UV exposure. As cells die and contract, an actin ring simultaneously forms and contracts to squeeze the cell out of the monolayer (Figure 2G,H, Movie S2). No gaps appear in the monolayer throughout the entire extrusion process, even when as many as five cells extrude within a field of view (Figure 2G). Thus, apoptotic cells extrude from endothelial layers using a similar process seen when epithelial cells are eliminated.

**Figure 2:**
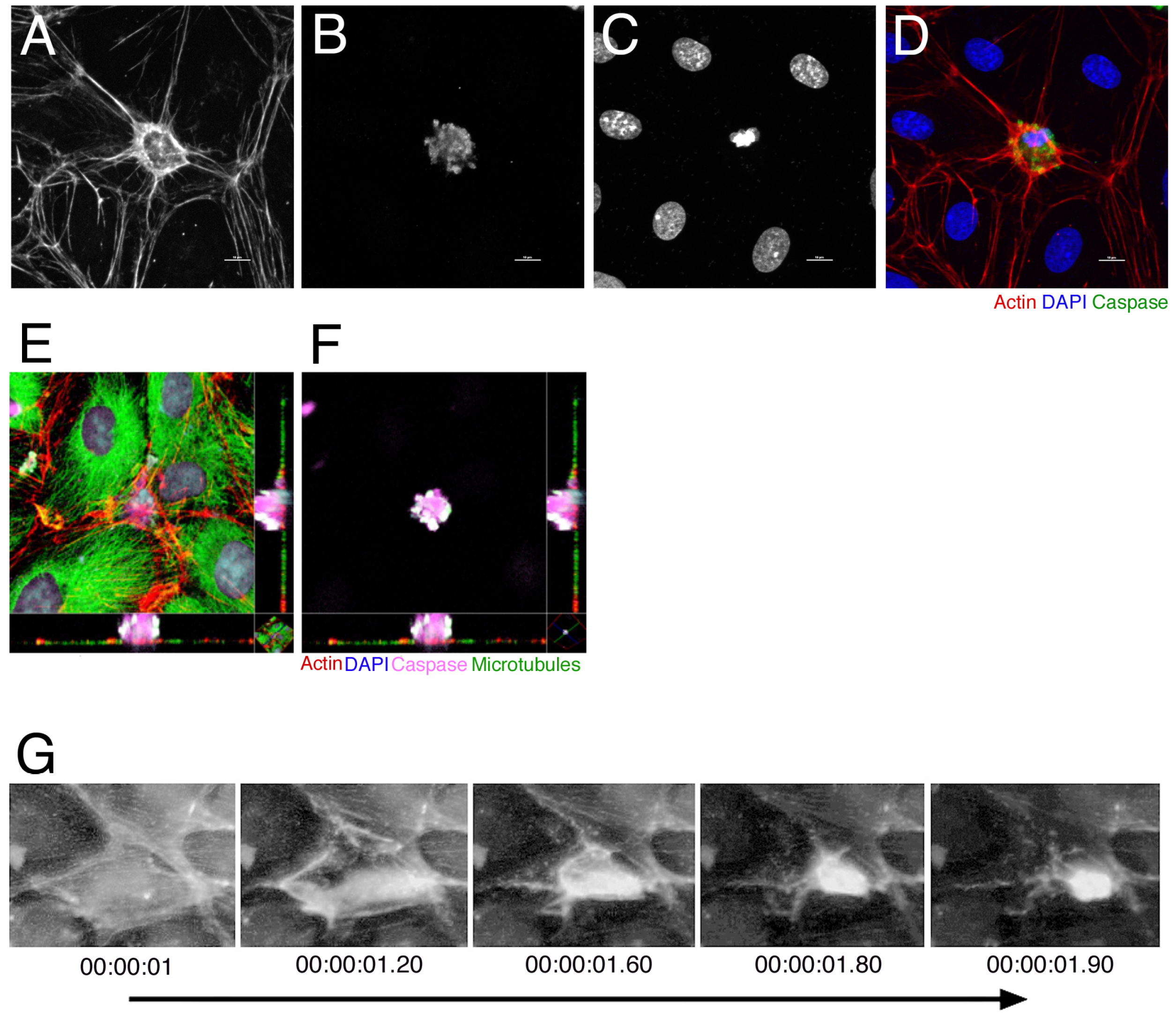
Endothelial cells extrude by intercellular actin ring contraction. A-D) Fluorescent micrographs of an endothelial cell undergoing extrusion. The actin ring, labeled with phalloidin (A), closes around a dying cell, as indicated by both active caspase staining (B) and condensed nuclei (C, DAPI). The final panel is a merged image with phalloidin in red, caspase in green, and nuclei in blue. Scale bar=10 E,F) Fluorescent confocal micrograph taken in the actin plane (E) and the caspase plane (F) with xy projections. G) Still images taken from a timelapse recording of LifeAct expressing endothelial cell extruding (Movie S2).

### Barrier function is preserved by apoptotic extrusion

To functionally test if the barrier is preserved during endothelial extrusion, we measured electrical resistance across HUVEC monolayers grown on filters following induction of apoptosis. Untreated HUVEC monolayers grown in an Electric Cell-substrate Impedance Sensing (ECIS) apparatus maintain high electrical resistance for up to six hours (Figure H&I). When exposed to UV^254^, HUVEC monolayers also maintained high electrical impedance over a six-hour period, similar to that seen by live intact endothelial cells (Figure 3H, I), even by the final time point when as many as 13% of the cells had died. Using epithelial cells as a control, we observe similar the same behavior of barrier preservation as that seen in endothelia (Rosenblatt *et al.*, 2001). Disruption of endothelial cell-cell contacts by treatment with either the calcium chelator EGTA or an antibody against V-cadherin to disrupt cadherin interactions resulted in a sharp decrease in resistance (Figure 3G).

**Figure 3:**
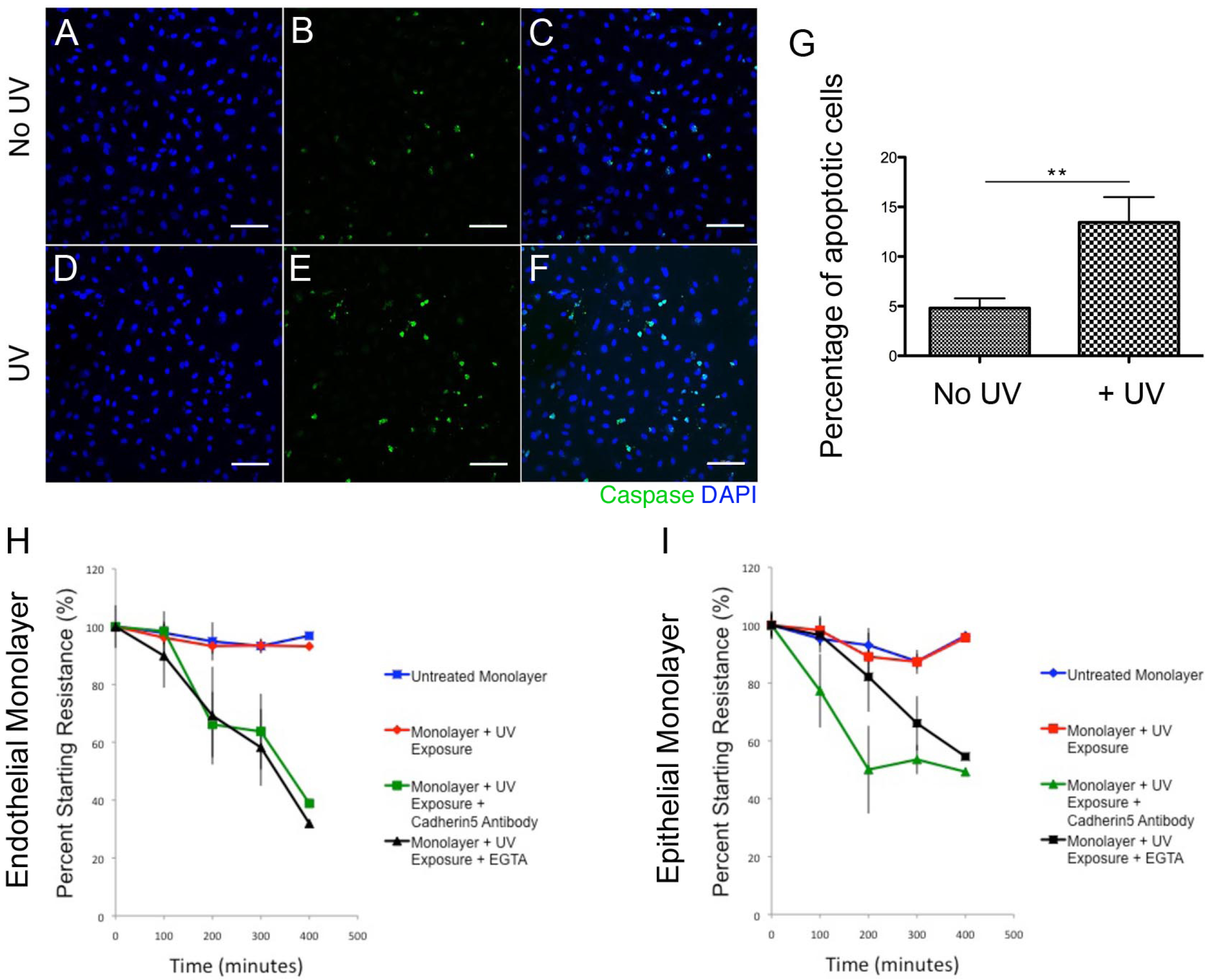
Endothelial barrier function is maintained when high rates of apoptosis are induced. A-F) Fluorescent images of extrusion in an endothelial monolayer. As indicated by nuclear (DAPI, blue; A,D) and caspase (green; B,E) staining. (C,F) are a merged image. I would like to see the actin here, too, to see if there are any non-apoptotic cell extrusions without UV. Scale bar=10μm G) Quantification of apoptosis, as determined by numbers of caspase-postive cells. H) Trans-endothelial measurement of resistance, an indicator of barrier function. UV-exposed endothelial cells are able to maintain resistance. Cadherin-5, functional blocking antibody, was used to control for junction disruption. EGTA was used as a positive control. I) The epithelial cell-line, MDCK, displays similar patterns of resistance to that seen in endothelial cells.

### Endothelial extrusion requires S1P-S1P_2_ signaling

An epithelial cell destined to extrude produces the lipid, S1P, which binds the S1P_2_ receptor in neighboring cells to trigger p115 RhoGEF-dependent activation of Rho that promotes extrusion (Slattum *et al.*, 2009; Gu *et al.*, 2011). Rho-activation triggers actin to polymerize and contract in both the dying cell and neighboring cells to squeeze the dying cell out. Blocking any step in this pathway inhibits extrusion, resulting in epithelial gaps at sites where cells die (Rosenblatt *et al.*, 2001, Gu *et al.*, 2011). To test if endothelia require the same S1P-S1P_2_-Rho pathway during extrusion, we used inhibitors to block different steps in this pathway when HUVEC endothelial monolayers were induced to die by UV^254^. Preventing S1P synthesis with a sphingosine-1-kinase inhibitor or inhibiting S1P activation of S1P_2_ with the S1P_2_ specific antagonist JTE-013 blocked 60% of apoptotic cells from extruding (Figure 4A). Uncontracted actin rings around caspase-3-positive cells, where the DNA and caspase-3 of the dying cell lies in the same plane as the surrounding endothelial cells indicate defective extrusion (Figure 4D,E). Additionally, inhibiting Rho kinase (ROCK) downstream of Rho with Y27632 blocked 90% of apoptotic cells from extruding in both non-treated and UV^254^-treated HUVEC monolayers (Figure 4B, F). These data demonstrate that endothelial extrusion requires the same S1P-S1P_2_-RhoA pathway identified for epithelial cell extrusion.

**Figure 4:**
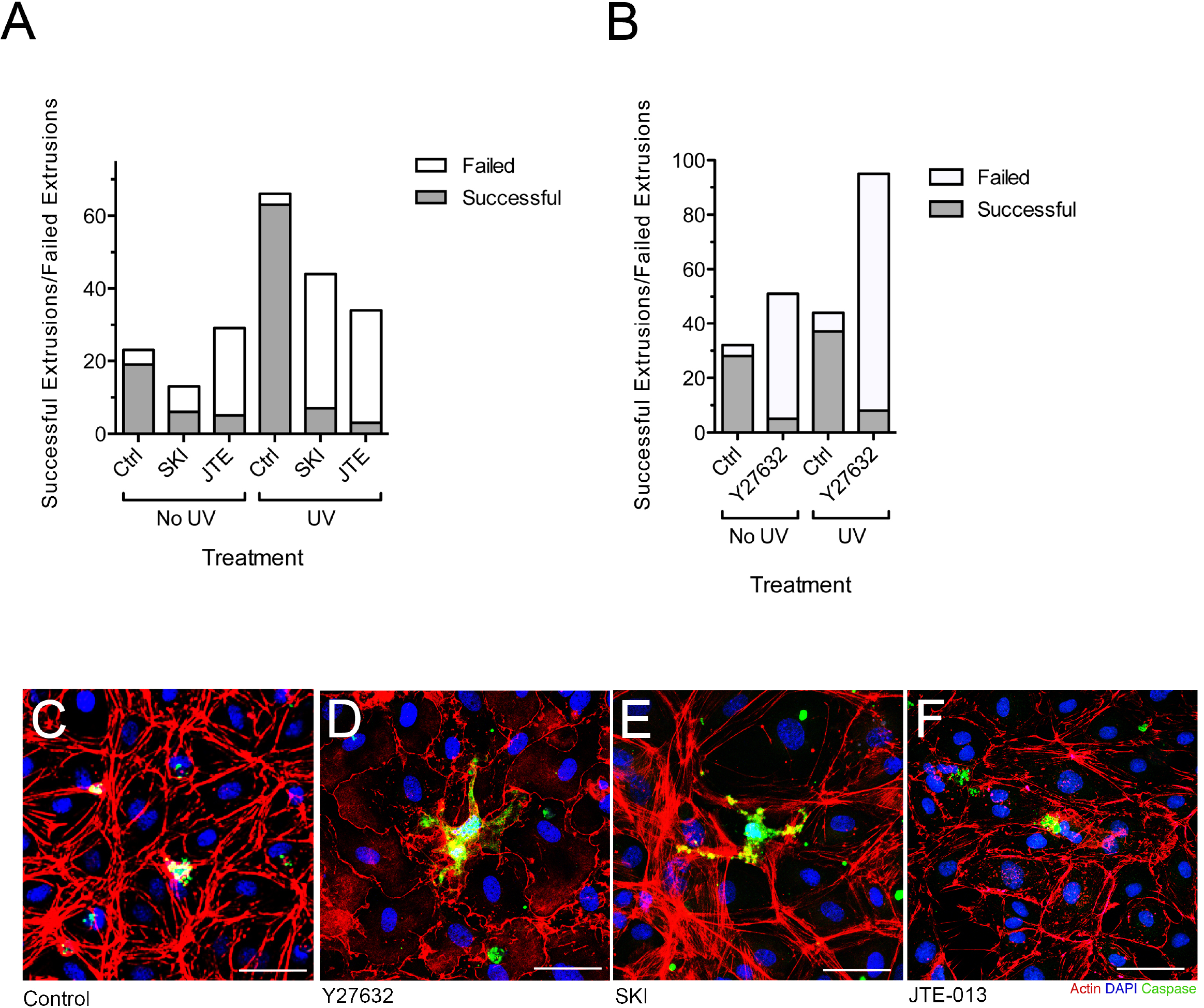
Endothelial extrusion uses the S1P-S1P2-Rho pathway. A) Quantification of sphingosine kinase (30μM) or JTE-013 inhibitor treated monolayers. B) Quantification of Y27632 (50μM) treated monolayers. Data in A and B was analyzed using a 3×2 contingency table and statistical significance measured by a Fisher′s Exact test. C-F) Cells treated with DMSO, Y-27632, sphingosine kinase inhibitor, or S1P2 antagonist (JTE-013) and immunostained for actin (red), active caspase-3 (green), and nuclei (DAPI, blue). Note that only DMSO extrudes by extruded efficient actin ring closure.

## Discussion

Our work demonstrates that endothelia extrude apoptotic cells to maintain a constant barrier when cells either are targeted to die or die spontaneously. Endothelia and epithelia appear to use similar cellular and signaling mechanisms to remove cells by extrusion. Our results suggest that diseases of vascular leakage may arise if the process of endothelial extrusion becomes defective.

The results we present define endothelial cell extrusion and the signaling that controls it may contribute to a better understanding of how vascular leakage diseases and blood borne diseases arise. Defects in extrusion could lead to hemorrhages in capillaries that are not lined with smooth muscle. In addition to causing leakage, disruption of endothelia can interfere with endothelial function as an anticoagulant surface and its ability to regulate thrombosis, which could result in excessive blood clotting (Deanfield *et al.*, 2007). Finally, since we have also discovered that disrupting epithelial extrusion can lead to cell masses and aberrant basal extrusion [Gu *et al.* 2015; Slattum *et al.*, 2014), disruption of endothelial extrusion could lead to aberrant blood vessel formation, such as angiogenic tangles or aberrant budding of blood stem cells and leukemias. Further study is required to determine if the concepts revealed by epithelial cell extrusion also relate to endothelial cell extrusion and whether misregulation of endothelial extrusion leads to vascular diseases.

## Materials and Methods

### Zebrafish

Housing and experiments were conducted according to IACUC protocol. The zebrafish line kdrl:GFP was a gift from Brent Bisgrove. 20-22hpf embryos were anesthetized with tricaine and 0.02mM phenylthiourea (PTU) and filmed in 1% low-melt agarose for 8 hours.

### Cell Culture

Human Dermal Microvascular Endothelial Cells (HMVEC-d, Lonza CC-2543) and Human Umbilical Vein Cells (HUVEC, Lonza C2519A) were cultured according to manufacturer’s instructions. Madin Darby Canine Kidney Cells (MDCK II) and HEK293T cells were cultured in DMEM high glucose with 5% FBS and 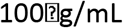 penicillin/streptomycin (all from Invitrogen) according to manufacturer’s instructions. LifeAct was expressed in HUVEC cells by nucleofection according to manufacturer’s protocol (Lonza).

### Immunofluorescent Cell Staining

Cells were grown to confluence on fibronectin coated glass cover slips and fixed using 4% PFA in PBS at 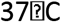 for 20 minutes. Cells were stained, as previously described [9], using a 1:100 dilution Alexa Fluor568 Phalloidin (Life Technologies A12380), a 1:1000 dilution of rabbit-anti-active Caspase-3 (BD Pharmingen 559565), washed and incubated with a 1:100 dilution of Alexa Fluor488 anti-rabbit (Life Technologies A11034). Cover slips were mounted using Prolong Gold Antifade Mountant with DAPI (Life Technologies P36930).

### Imaging

Fluorescent micrographs were taken using a Nikon A1R confocal microscope. Visual quantification of cells was done using a Leica TCS SP5 microscope. Live cell time-lapse imaging was done using an Olympus IX81 automated microscope. Live imaging and IR ablations in zebrafish were done using a Prairie two-photon microscope.

### Barrier Function Assay

Cells were grown to confluence on a fibronectin-coated ECIS 8W10E+ gold electrode chamber slide (Biophysics). Resistance across the monolayer was measured using the Electrical Cell-substrate Impedance System (ECIS) from Applied Biophysics. Cells were allowed to rest for 30 minutes prior to any treatment. Values were normalized against control and quantified.

### Apoptosis Induction and Drug Inhibitors

To induce apoptosis, cells were irradiated at 400 in a UV Stratalinker 1800. Inhibitors were used at the following concentrations prior to UV exposure: 200 μM Y27632 dihydrochloride (Rho kinase inhibitor, Tocris 1254), 10 μM S1P2 antagonist JTE-013 (Tocris 2392), or 30 μM SKI II (EMD 509106).

## Supplemental Figure legends

**Movie S1:**
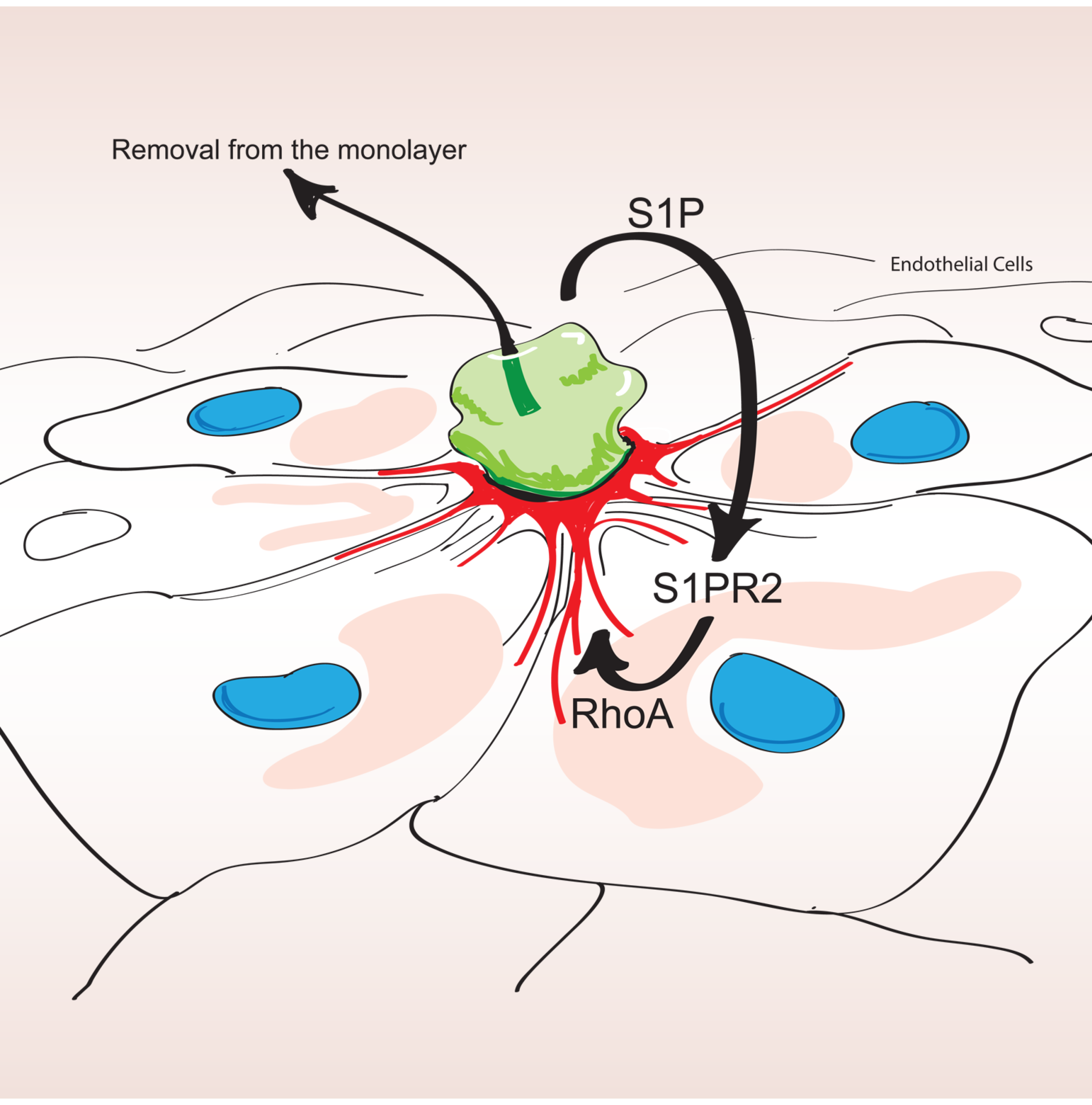
Endothelia extrude dying cells *in vivo.* Timelapse of kdrl:GFP zebrafish CCV treated with 720nm two-photon laser. Images were taken over an eleven hour period with extrusions beginning to occur approximately four hours into filming. In this movie, two events can be observed. Corresponding still images can be seen in Figure 1B.

**Movie S2:**
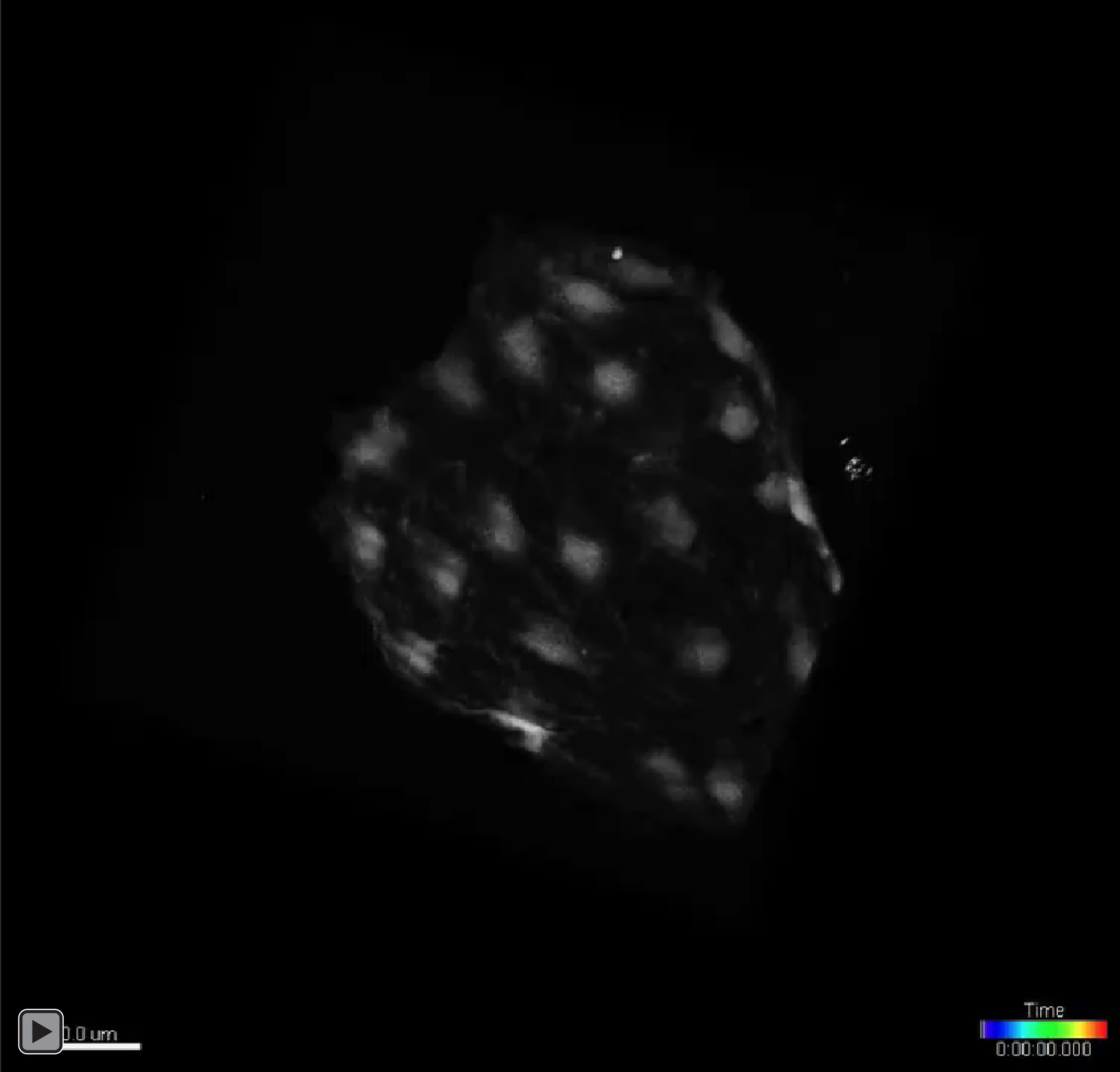
Endothelial cells extrude by intercellular actin ring contraction. Timelapse of LifeAct labeled HUVEC cells. Images were taken over an eight hour period with extrusion events beginning to occur approximately four to five hours into filming. In this movie, five events can be observed. Corresponding still images can be seen in Figure 2G.

